# Biases and Blind-Spots in Genome-Wide CRISPR Knockout Screens

**DOI:** 10.1101/2020.01.16.909606

**Authors:** Merve Dede, Eiru Kim, Traver Hart

**Affiliations:** Department of Bioinformatics and Computational Biology, The University of Texas MD Anderson Cancer Center, Houston, Texas, USA; Graduate School of Biological Sciences, The University of Texas MD Anderson Cancer Center, Houston, Texas, USA; Department of Cancer Biology, The University of Texas MD Anderson Cancer Center, Houston, Texas, USA

## Abstract

It is widely accepted that pooled library CRISPR knockout screens offer greater sensitivity and specificity than prior technologies in detecting genes whose disruption leads to fitness defects, a critical step in identifying candidate cancer targets. However, the assumption that CRISPR screens are saturating has been largely untested. Through integrated analysis of screen data in cancer cell lines generated by the Cancer Dependency Map, we show that a typical CRISPR screen has a ∼20% false negative rate, beyond library-specific false negatives previously described. Replicability falls sharply as gene expression decreases, while cancer subtype-specific genes within a tissue show distinct profiles compared to false negatives. Cumulative analyses across tissues suggest only a small number of lineage-specific essential genes and that these genes are highly enriched for transcription factors that define pathways of tissue differentiation. In addition, we show that half of all constitutively-expressed genes are never hits in any CRISPR screen, and that these never-essentials are highly enriched for paralogs. Together these observations strongly suggest that functional buffering masks single knockout phenotypes for a substantial number of genes, describing a major blind spot in CRISPR-based mammalian functional genomics approaches.

## Introduction

The search for essential genes - genes whose loss of function results in a severe fitness defect - has been of outstanding interest to the scientific community. Prior to advanced genomic technologies, the assumption was that the majority of genes were essential for life (Horowitz and Leupold, 1951). This idea was dismissed by several studies that utilized saturating random mutagenesis to show that in *C. elegans* and *S. cerevisiae*, 12-15% of the genome was estimated to be essential (Brenner, 1974; Goebl and Petes, 1986). These studies were limited by the methods at the time and the lack of the availability of complete genome sequences.

After improvements in shotgun sequencing, initial studies to define essential genes in bacteria were driven by the desire to identify antimicrobial targets with the first minimal genome screen performed in *Mycoplasma genitalium* (Hutchison et al., 1999). Later studies revealed the essential genes in other bacteria including *M. tuberculosis*, *P. aeruginosa* and *H. influenza* (Sassetti et al., 2001) (Jacobs et al., 2003)(Akerley et al., 2002).

With the advances in genome technologies that enabled sequencing of eukaryotic organisms, systematic gene knockout studies were performed in *S. cerevisiae*, identifying essential genes by deletion of open reading frames in the yeast genome (Giaever et al., 2002; Winzeler et al., 1999). This method identified that 17% of yeast genes were essential for growth in rich medium (Winzeler et al., 1999). However, a later study showed that a binary classification of genes into essential and non-essential was misleading due to the context dependent nature of gene essentiality and that 97% of yeast genes showed some growth phenotype under different environmental conditions (Hillenmeyer et al., 2008).

Developments in RNA interference (RNAi) technology provided valuable insight into detection of fitness genes. Generation of genome scale RNAi libraries to conduct genome-wide RNAi screens facilitated the study of essential genes in multiple organisms (Dietzl et al., 2007; Kamath et al., 2003; Meister and Tuschl, 2004; Moffat and Sabatini, 2006). In these RNAi screens, 30% of the genome was shown to be essential in *D.melanogaster* cell lines, (Dietzl et al., 2007), compared to only 8.5% of the *C.elegans* genome in whole worms (Kamath et al., 2003).

Identifying essential genes in human cancer cells is of special interest in oncology since the cancer-specific essential genes represent genomic vulnerabilities that can potentially be targeted with novel therapeutic agents. An initial study showed that some colorectal cell lines were dependent on a specific KRAS mutation for growth and survival (Shirasawa et al., 1993). Later this idea was explored under the term “oncogene addiction” that describes the dependency of cancer cells on specific oncogenes for sustained growth and proliferation (Weinstein and Joe, 2008). To identify these oncogenes, RNAi screens were performed on small arrays of cancer cells to search for essential genes (Moffat et al., 2006; Schlabach et al., 2008; Silva et al., 2008). Subsequent larger-scale efforts such as the Project Achilles of the Broad Institute focused on context specific gene essentiality across 216 human cancer cell lines screened with an shRNA library (Cowley et al., 2014). Similarly, another relatively big scale study in 72 cell lines (Marcotte et al., 2012) produced consistent results with the previous studies, indicating confidence in the methodology. Even though significant efforts have been made to optimize reagent design and analytical methods, RNAi technology was shown to have significant limitations such as incomplete loss of function due to RNAi, decreased sensitivity for genes with low expression levels (false negatives) and confounding off-target effects (false positives) (Boutros and Ahringer, 2008; Echeverri et al., 2006; Hart et al., 2014).

More recently, adaptation of the bacterial CRISPR-Cas9 system to mammalian cells enabled genome-scale approaches to define human essential genes. Studies using this technology revealed that mammalian cells have more essential genes than RNAi screens were able to detect and that, at the same false discovery rate, CRISPR screens generated 3-4 times more essential genes (Hart et al., 2014). Moreover, multiple groups revealed lists of ∼2000 highly concordant human essential genes, and comparison of CRISPR technology to orthogonal techniques such as random insertion of gene traps also showed consistent results (Blomen et al., 2015; Hart et al., 2015; Wang et al., 2015). These findings were initially thought to indicate that the CRISPR-

Cas9 screens are saturating and that a well-designed screen can detect a cell’s full complement of essential genes. However, it is still poorly understood how the possible systematic biases and blind spots in CRISPR screens affect our understanding of human gene essentiality. In the absence of a ground truth, the actual true positive, false positive and false negative rates are in an average genome-wide CRISPR-Cas9 knockout screen are still unknown.

Moreover, even with the CRISPR technology, the number of essential genes detected through these screens is still far less than the number of genes expressed in a given cell line. Large-scale experiments exploring genetic and environmental buffering in both yeast (Costanzo et al., 2010, 2016; Hillenmeyer et al., 2008; VanderSluis et al., 2014) and C. elegans (Ramani et al., 2012) suggest that virtually every gene is required for optimal growth in some condition. A major open question remains whether these findings hold true for human cells generally and cancer cells specifically.

In this study, we examine some of the biases and blind spots characteristic of genome-wide CRISPR-Cas9 knockout screening. Using publicly available genome wide screen data from 517 genetically heterogeneous cell lines from the Cancer Dependency Map initiative (Meyers et al., 2017; Tsherniak et al., 2017), we demonstrate the systematic biases in these screens, investigate and model the actual number of essential genes identifiable with CRISPR technology, estimate the false discovery rate (FDR) and false negative rate (FNR) in a “typical” CRISPR screen, and reveal blind spots that can offer fruitful areas for further research.

## Results

To systematically evaluate the biases and blind spots in genome-wide CRISPR-Cas9 knockout screening, we processed the raw read counts of loss of function screens performed in 517 genetically heterogeneous cell lines from the 2018Q4 release of the publicly available Avana data (Meyers et al., 2017). We applied our computational pipeline as described in the methods section to correct for copy number effects using the previously described CRISPRcleanR algorithm (Iorio et al., 2018) and assign an essentiality score (Bayes Factor, BF) to each gene in every screen using an updated version of our BAGEL algorithm (Hart and Moffat, 2016). After applying quality control metrics (see Methods), our final dataset included 446 screens meeting the F-measure criteria of 0.80 and above (Supplementary Figure 1A-B, Supplementary Table1). These 446 cell lines are derived from 23 tissue types, with varying representation (Figure 1A).

**Figure 1.**
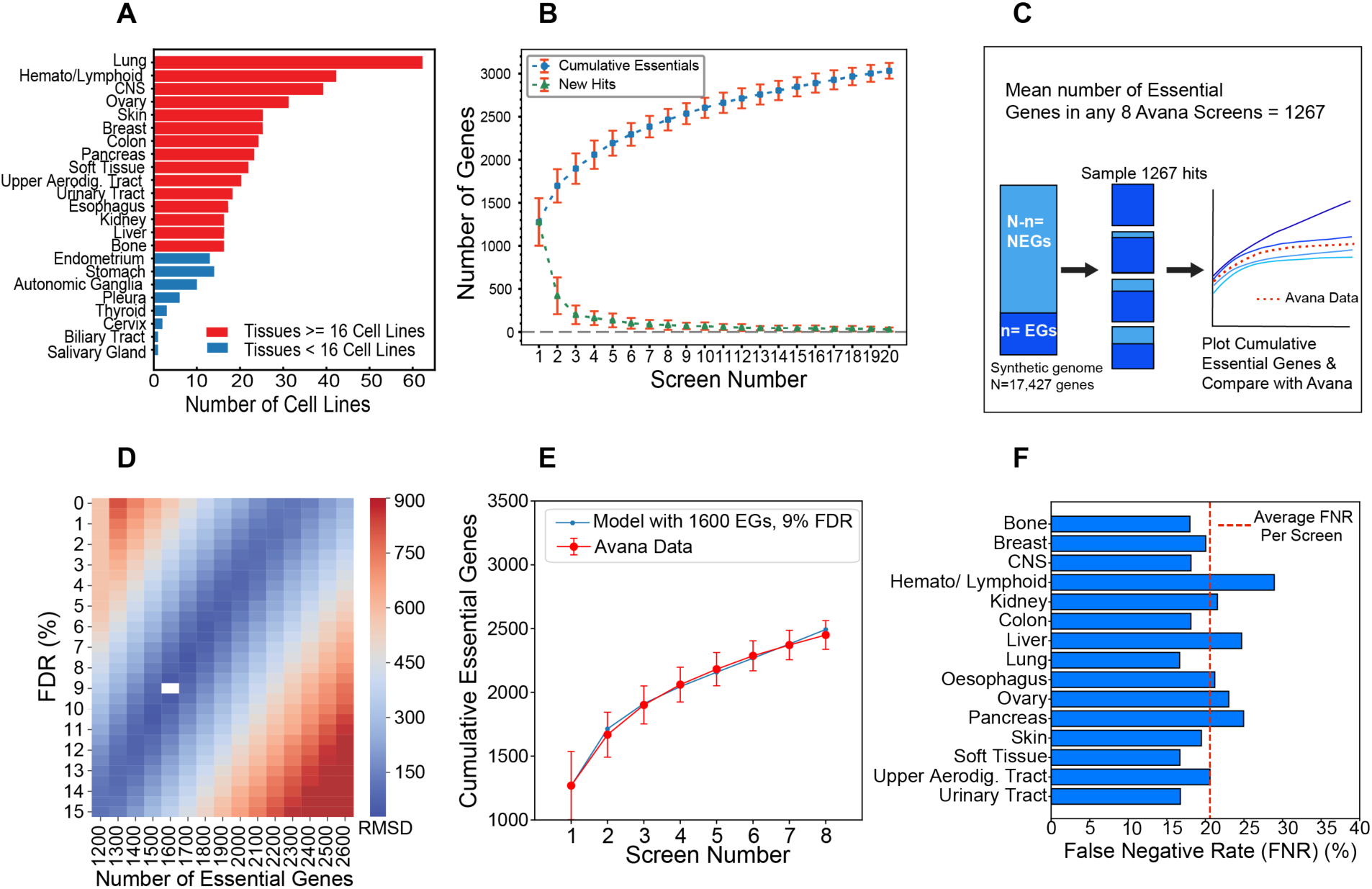
Synthetic genome modeling of Avana data. **A)** The representation of each tissue type in the 446 high-performing screens in the Avana data. Tissue types with >=16 high-performing cell lines are indicated with red bars. **B)** The cumulative number of essential genes (BF>=10) and newly discovered essential gene curves in any randomly selected set of 20 Avana screens show a curve that converges into a positive slope. **C)** Diagram overview of the synthetic genome modeling of essential genes. From a synthetic genome of 17,427 genes, 1267 hits were randomly sampled from the essential (n) and non-essential gene (NEG) populations based on the defined false discovery rate (FDR) in the simulation. The resulting cumulative essential hits across 8 iterations were plotted and compared to the mean cumulative essentials curve from the Avana data. **D)** The synthetic genome modeling revealed that the best fitting model was the one with n=1600 genes and 9% FDR. **E)** Plot showing the model with the best fit (in blue) to the Avana data (red). **F)** An average screen false negative rate (FNR) was determined by comparing the number of essential genes from the best fit model in each tissue type to the mean number of observed essentials in each tissue type. The red dashed line indicates a mean FNR of 20.6% across all tissue types tested.

### Synthetic genome modeling of essential genes in 446 screens

To estimate the total number of essential genes in a cell, we first considered an approach based on the cumulative observations across all screens. The expectation is that, for a sufficient number of identical screens with no false positives, a plot of the cumulative number of essential genes would flatten to zero slope as the total population of cell-essential genes was identified. In contrast, in screens with either cellular heterogeneity or some low false discovery rate (or both), the slope of the cumulative essential plot would remain positive, reflecting the ongoing accumulation of false positives (or, alternatively, cell-specific essential genes) in otherwise saturated screens. We previously applied this principle to estimate the total population of essential genes assayable by shRNA screens (Hart et al., 2014).

We plotted the cumulative essential genes across sets of 20 cell lines randomly selected without replacement from all screens (100 iterations). Filtering for genes with BF>10 in each screen, a strict threshold representing a posterior probability of gene essentiality of ∼99% (Supplementary Figure 1C; see Methods), yielded a curve that converged to a positive slope (Figure 1B), similar to that shown previously in shRNA screens. We reasoned that this curve represents three factors: first, there exists a fixed population of essential genes across all screens. Second, the screening platform does not reliably capture all of these genes in a single experiment. Each screen therefore carries some unknown false negative rate, and multiple screens are required before saturation. Third, after saturation, additional screens continue to detect some combination of false positives and context-specific essential genes that were either not detectable or not present in the prior set of screens, and that the rate at which these genes are observed offers some estimate of the false discovery rate of each screen.

To model these factors, we carried out repeated screens *in silico* and compared synthetic cumulative essential curves to those derived from the data. Starting with a genome of N=17,427 genes – the number of genes tested in the Avana library – we arbitrarily defined *n* essential genes, leaving N-*n* nonessential. We further defined an arbitrary screen false discovery rate between 1-15%. Then we repeatedly sampled this genome with a screen that randomly drew 1,267 hits – the mean number of hits at BF>10 across all Avana screens -- from the essential and nonessential populations based on the defined FDR (e.g. at 10% FDR, 127 nonessentials and 1,140 essentials were randomly selected; see Figure 1C). Finally, we determined the cumulative hits across eight iterations, estimating that eight samples was a good estimate of screen saturation in the data (Figure 1B) and judging that it was more important to fit the model to our observations in this region than in the saturated region. We calculated the root-mean-squared deviation from the mean cumulative essentials curve determined from the Avana data and plotted RMSD vs. the two parameters of the model (Figure 1D), observing that the best fit occurred with *n*=1,600 essential genes and FDR=9% (Figure 1E). Notably, a region of good fits, with RMSD < 2xRMSD_min_, occurs between *n=*1,400-1,900 essential genes and a corresponding decrease in per-screen FDR from 9% to ∼5% (Supplementary Figure 1D).

Given the broad range of lineages from whence the screened cell line models were derived, it seems clear that context-specific essential genes will be included in these putative false positives. To minimize the contribution of these tissue-specific essentials, we repeated the analysis using screens derived only from a single tissue or lineage, filtering for lineages represented by at least 16 high-quality screens (Figure 1A, red; n=15 lineages). Every tissue yielded a similar cumulative essential curve (Supplementary Figure 2). We repeated the synthetic genome modeling approach in each tissue, with remarkably similar results (Supplementary Figures 3 and 4). By comparing the best-fit number of essential genes to the mean number of hits observed each screen, we calculated an average false negative rate across each tissue (Figure 1F). Across all tissues, we determined the mean FNR to be ∼20% in each screen.

### Saturation modeling to differentiate essential genes and false positives

While the synthetic genome modeling approach described above can estimate the total number of essential genes in a tissue, it does not provide any way to differentiate true hits from false positives. To address this issue, we took an alternative view of the saturating behavior of CRISPR screens. Based on our judgment that screening in virtually all lineages achieved saturation after roughly eight cell lines had been effectively screened, we again selected the lineages with at least twice this number of cell lines (Figure 1A). From each lineage, we randomly selected eight screens (“initial screens”) without replacement and determined the number of cell lines in which each gene was classified as essential (BF>10; Figure 2A). We then randomly selected an additional eight screens (“subsequent screens”), again without replacement, and determined the number of cell lines in which each gene was classified as new hits – that is, BF>10 but not a hit in any of the initial eight screens. We assumed that all of these new hits were false positives, and that the histogram of observations of these false positives estimates the frequency of false positives in the initial screens. It is almost certainly not the case that all of these are actually false positives, given the known presence of tumor subtypes within each tissue/lineage, the high likelihood of subtype-specific essential genes, and the probability that any given subtype escaped being selected in the initial eight screens. However, this assumption is useful for modeling purposes, as it provides an estimate of the upper bound of the false discovery rate using this saturation modeling approach. We repeated this process 100 times and plot the resulting histogram in Figure 2A.

**Figure 2.**
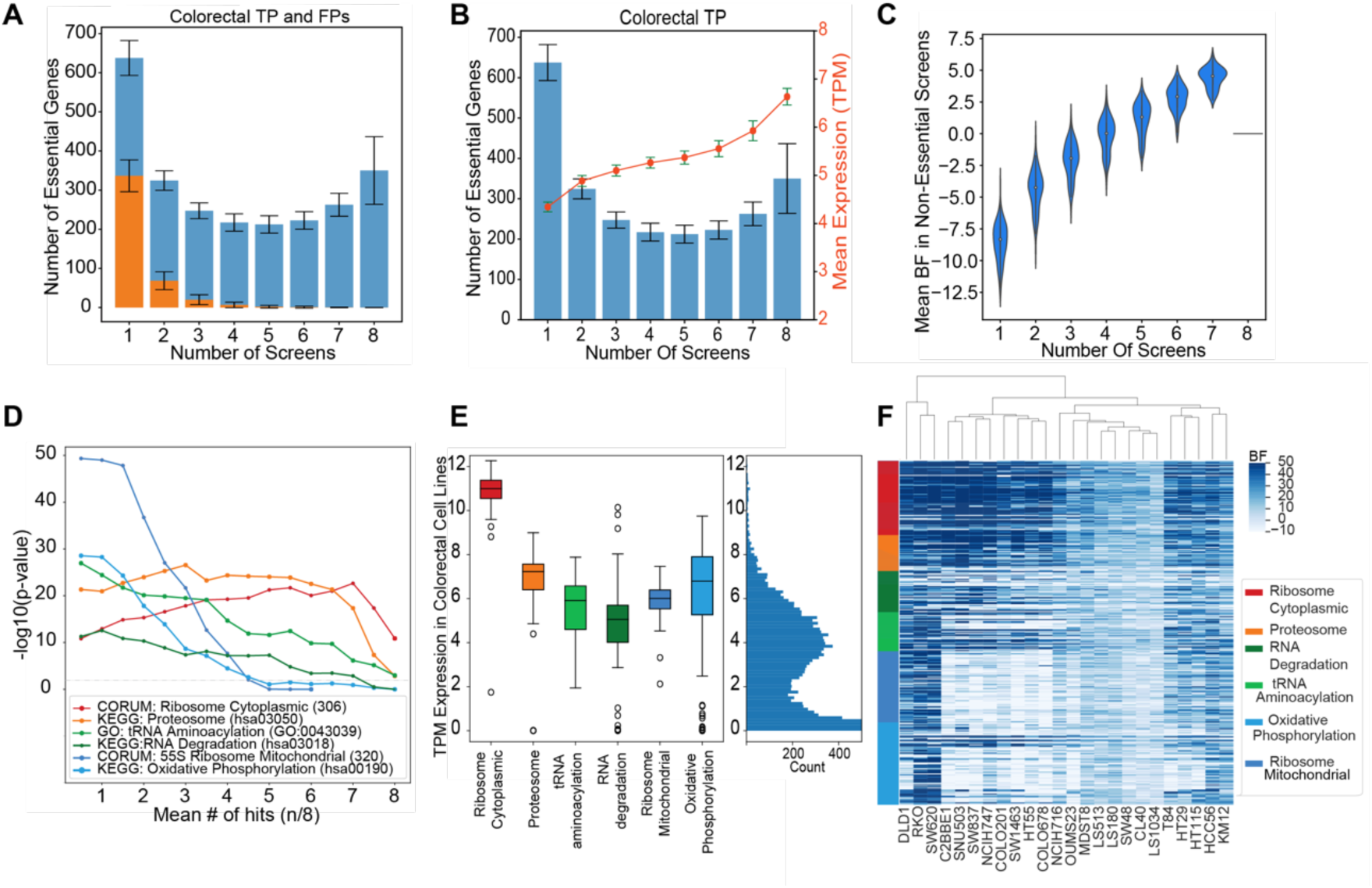
Saturating modeling approach, differentiating essential genes, false positives, false negatives and subtype-specific essential genes. **A)** Histogram showing the distribution of the number of essential genes and the number of cell lines in which each gene was classified as essential in colorectal cancer cell lines. Blue bars represent the distribution of true positives (TPs) and the orange bars represent the frequency of false positives (FPs). The error bars represent the standard deviation of observed essentials in 100 iterations. **B)** For the TPs in colorectal cancer cell lines depicted in **A**, the mean mRNA expression (log2(TPM)) of the gens in each bin shows higher expression where more frequently observed essential genes show higher levels of expression. **C)** Violin plot showing the distribution of the essentiality scores (Bayes Factor, BFs) of the TP genes in **A and B**, in the screens where they were not observed as essential. **D)** Functional enrichment of genes in colorectal cancer cell lines based on their mean number of hits observations out of 8 screens in 100 iterations. Grey dashed line indicates significance at p=0.01 **E)** mRNA expression (log2(TPM)) in colorectal cancer cell lines of the genes involved in the enriched pathways shown in **D.** Marginal histogram on the right shows the distribution of expression levels of all genes assayed in the Avana library. **F)** Hierarchical clustering of the colorectal cancer cell lines based on the essentiality scores of the genes involved in pathways showed in **D and E**.

Notably, the putative false positives follow the expected distribution: most are detected in only a single screen (Figure 2A). We can estimate both binwise and cumulative false discovery rates (Supplementary Table 2) by comparing the ratio of putative false positives per bin to the total number of hits. We assess that genes observed in 3 or more of 8 randomly selected screens represent the high-confidence set of essential genes in a given lineage, including both genes that are frequently false negatives as well as those that are subtype-specific within a lineage.

### False negatives vs. subtype-specific genes

The familiar U-shaped histogram in Figure 2A carries strong implications for the underlying experiment. Genes observed in an intermediate number of screens (3 to 6, out of 8) are either false positives that are repeatedly observed, false negatives that are repeatedly missed, or subtype-specific genes that are only hits in some cells, violating the modeling assumption that the cells are identical (in reality, some combination of the three is likely). We show from the hit frequency in subsequent screens that these genes are unlikely to be false positives. Here we attempt to differentiate between false negatives and context-dependent genes.

First, we find the mean mRNA expression of genes in each bin (Figure 2B). A clear trend emerges, whereby more frequently observed hits show higher gene expression. Genes observed in only one of eight screens, highly enriched for false positives, show markedly lower expression. In addition, putative false positives from subsequent screens show a similarly lower average gene expression than more frequently observed hits (Supplementary Figure 5). Second, we examine the BF scores of genes in screens where the gene is not a hit. This measures whether a gene that is essential (BF>10) in, for example, 5 screens is truly nonessential in the remaining 3 screens (BF<-10) or falls in the indeterminate range near BF=0. Figure 2C shows that, the more frequently a gene is classified as a hit in the initial screens, the higher its average BF in screens where it is not a hit. This is strongly consistent with false negatives rather than context-specific hits.

Finally, we measured functional enrichment of gene annotations as a function of stringency thresholds. We plotted the P-value of annotation enrichment for several terms that demonstrate the major trends in the data. Genes associated with the cytoplasmic ribosome, which should be essential in every cell, show peak enrichment at high hit frequency (hits in n>=7 of 8 screens in 100 random samples; Figure 2D). Consistent with the expression bias shown in Figure 2B, these genes are very highly expressed in the cell (Figure 2E). Similarly, genes encoding proteasome subunits are critical for proliferation of all cells, show near-maximal enrichment at high frequency of observation (n>6.5, Figure 2D), and are relatively highly expressed (Figure 2E). In contrast, genes involved in tRNA aminoacylation and RNA degradation—which should also, in principle, be universally essential—show consistent increase in enrichment as frequency of observation is relaxed (Figure 2D), and these genes are expressed at intermediate levels (Figure 2E). Taken together, these trends are consistent with a significant false negative rate among moderately expressed genes that should otherwise exhibit consistent fitness defects across cell lines. Moreover, this trend is easily differentiated from context-specific modular functions: genes related to mitochondrial translation and oxidative phosphorylation only show enrichment at low frequency of observation (Figure 2D), despite their robust gene expression (Figure 2E). A summary of gene essentiality scores and trends in the cell lines displayed here is shown in Figure 2F, with context-dependent oxphos genes driving the hierarchical clustering of cell lines. A complete table of gene frequency of observation by tissue type is presented in Supplementary Table 3.

### Context-specific essential genes define lineage relationships

After judging that genes observed in 3 or more (of 8) screens represent the high-confidence set of essential genes in a given lineage, we identified 954 genes that are hits at that frequency in all 15 lineages we evaluated (Figure 3A). In addition, each lineage carries an additional 300-600 context-specific essential genes (Figure 3A, inset). These additional genes are also widely, but not universally, shared across backgrounds: each lineage has only three (CNS) to 49 (hematopoietic) genes uniquely essential to that lineage (Figure 3A). Many known gene-tissue relationships are described in this set of unique context essentials. For example, the *SOX10* transcription factor was found to be essential in only skin cells where it plays a major role in the production and function of melanocytes (Harris et al., 2010; Nonaka et al., 2008). *CTNNB1* and *TCF7L2* are essential only in colorectal cancer cell lines, where activation of the Wnt pathway results in accumulation of *B-*catenin that interacts with and acts as a coactivator for *TCF7L2* that in turn activates downstream genes responsible for colorectal cancer cell survival as well as resistance to chemo-radiotherapy (Albuquerque and Pebre Pereira, 2018; Emons et al., 2017; Murphy et al., 2016). ER+ breast cancer cell lines specifically depend on transcription factors *FOXA1* and *GATA3,* which are overexpressed in ER+ breast carcinomas (Albergaria et al., 2009; Davis et al., 2016). *E2F1*, which was uniquely essential in only pancreatic cancer cells, is known to regulate both pancreatic B cell development and cancer growth by increasing the expression of *PDK1* and *PDK3* which results in increased aerobic glycolysis and growth in pancreatic cancers (Denechaud et al., 2017; Kim and Rane, 2011; Wang et al., 2016). Nevertheless, genes unique to a particular context are very rare, while hundreds of genes are shared across some but not all lineages. We tested whether clustering of these integrative tissue profiles of gene essentiality would recapitulate known lineage relationships. Hierarchical clustering of all context essential genes (Figure 3B) clearly separates epithelial-derived carcinomas from cancers of hematopoietic and bone/soft tissue origins. A complete table of common and context essential genes are listed in Supplementary Table 4.

**Figure 3.**
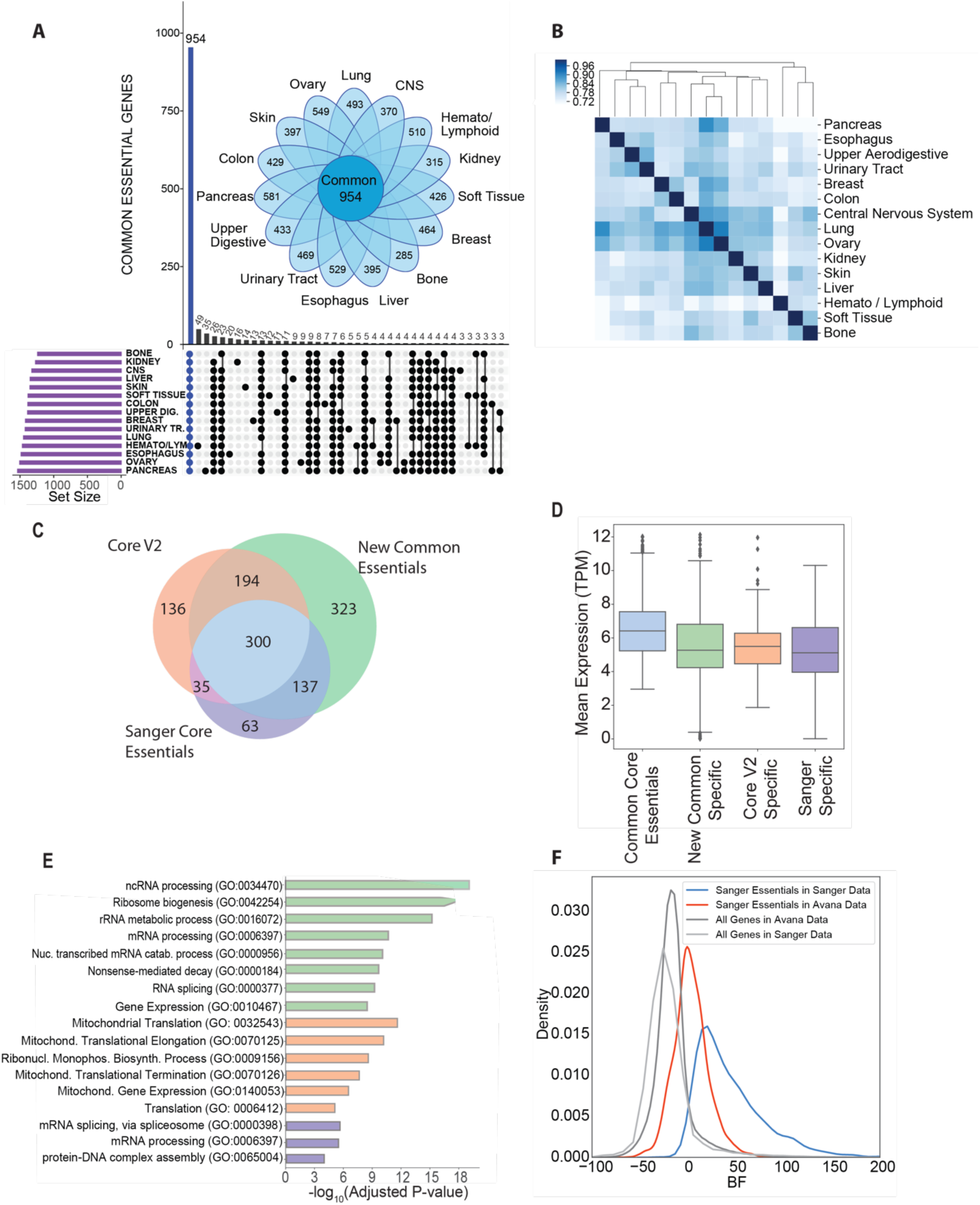
High-confidence essential genes, their characteristics and comparison to previously defined gold standard sets of core essential genes. **A)** Upset plot showing the number of intersection between high-confidence essential genes in each tissue type. Inset: Daisy plot showing the relationship between high-confidence context-essential genes and common essential genes. Genes essential in a given tissue type is represented by the petals of the daisy indicating their numbers for each tissue type. The petals overlap to varying degrees, but all tissues share the common set of essential genes (N=954). **B)** Hierarchical clustering of the high-confidence context-essential genes in different tissues based on their mean number of hits observations out of 8 screens in 100 iterations. **C)** Venn diagram comparing common essential genes to previously defined gold standard set of core essential genes. **D)** Box plots showing the mean mRNA (TPM) expression of common core essentials (n=300), genes unique to new common essentials identified here (n=323), Core V2 specific essentials (n=136) and core essential genes specific to the Sanger dataset. **E)** Biological process enrichment for core essential genes unique to a specific approach. **F)** Comparison of the distribution of essentiality scores of Sanger specific core essentials in common cell lines between the Avana and Sanger data.

### Comparing common essentials to previous gold standards

The common essentials defined here include genes that are identified as hits in every tissue at a frequency of at least 3/8 screens. They should, in principle, define a superset of previously defined gold standard sets of essential genes. We compared our set of 954 common essentials to the Core Essential Genes v2 that we previously defined as a gold standard training set for our BAGEL algorithm (Hart et al., 2017), as well as the core essentials recently published Sanger dataset derived from 342 CRISPR screens performed at the Wellcome Trust Sanger Centre. (Behan et al., 2019). Since the initially reported Avana data does not contain guides targeting genes on the X chromosome, we removed these genes from both other datasets to perform direct comparisons. Common essentials comprise 494 of 665 (74%) CEGv2 genes and 437 of 535 (82%) of Sanger core essentials (Figure 3C). The 300 genes common to all three approaches have median gene expression roughly twice that of the genes unique to each approach (median log2(TPM) 6.4 vs 5.3 for new common specific genes, 5.5 for CEGV2 and 5.1 for Sanger specific genes; Figure 3D), consistent with an increased false negative rate among essential genes with moderate levels of mRNA expression. This is also consistent with the increased overlap of common essentials with CEGv2 as stringency is increased (Supplementary Figure 6). Genes specific to each approach are listed in Supplementary Table 5.

Interestingly, the genes unique to each dataset show a strong bias reflecting the approach used. Genes unique to the common essentials defined here are highly enriched for ribosome genesis and mRNA processing genes (Figure 3E). Genes unique to CEGv2 are strongly biased toward nuclear genes encoding subunits of the mitochondrial ribosome and the electron transport chain. Cellular dependence on these biological processes is subtype-specific in the approach described here (Figure 2D,E,F) and these genes are likely excluded from the Sanger core essentials for the same reason. Sanger-specific core essentials do not show strong functional enrichment, compared to the other groups (Figure 3E). In fact, among the 63 Sanger-specific essentials, 14 were not targeted in the Avana library and the remainder tend to show intermediate BF scores in 117 high-performing Avana screens of the same cell lines (Figure 3F). This is consistent with there being a set of CRISPR library-specific false negatives, as previously reported by Kosuke Yusa and colleagues (Ong et al., 2017), which may be independent of the expression-associated false negatives reported here.

### Genetic buffering is systematically excluded from monogenic CRISPR screens

Beyond the false negatives detected by the saturation analyses described above, we sought to determine whether other systematic sources of bias exist in the CRISPR knockout screen data. We first attempted to correct for lineage-specific effects by defining a set of 7,378 constitutively expressed genes (logTPM > 2 and stdev < 1 across the cell lines for which Avana screening data was available; Supplementary Figure 7A). Common essentials are almost all constitutively expressed (Figure 4A), while roughly two-thirds of context-dependent essentials are also constitutively expressed (Figure 4A). Conversely, nearly half of all constitutively expressed genes (N=3,361; 46%) show no strong knockout phenotype in any Avana CRISPR screen (“never essentials”).

**Figure 4.**
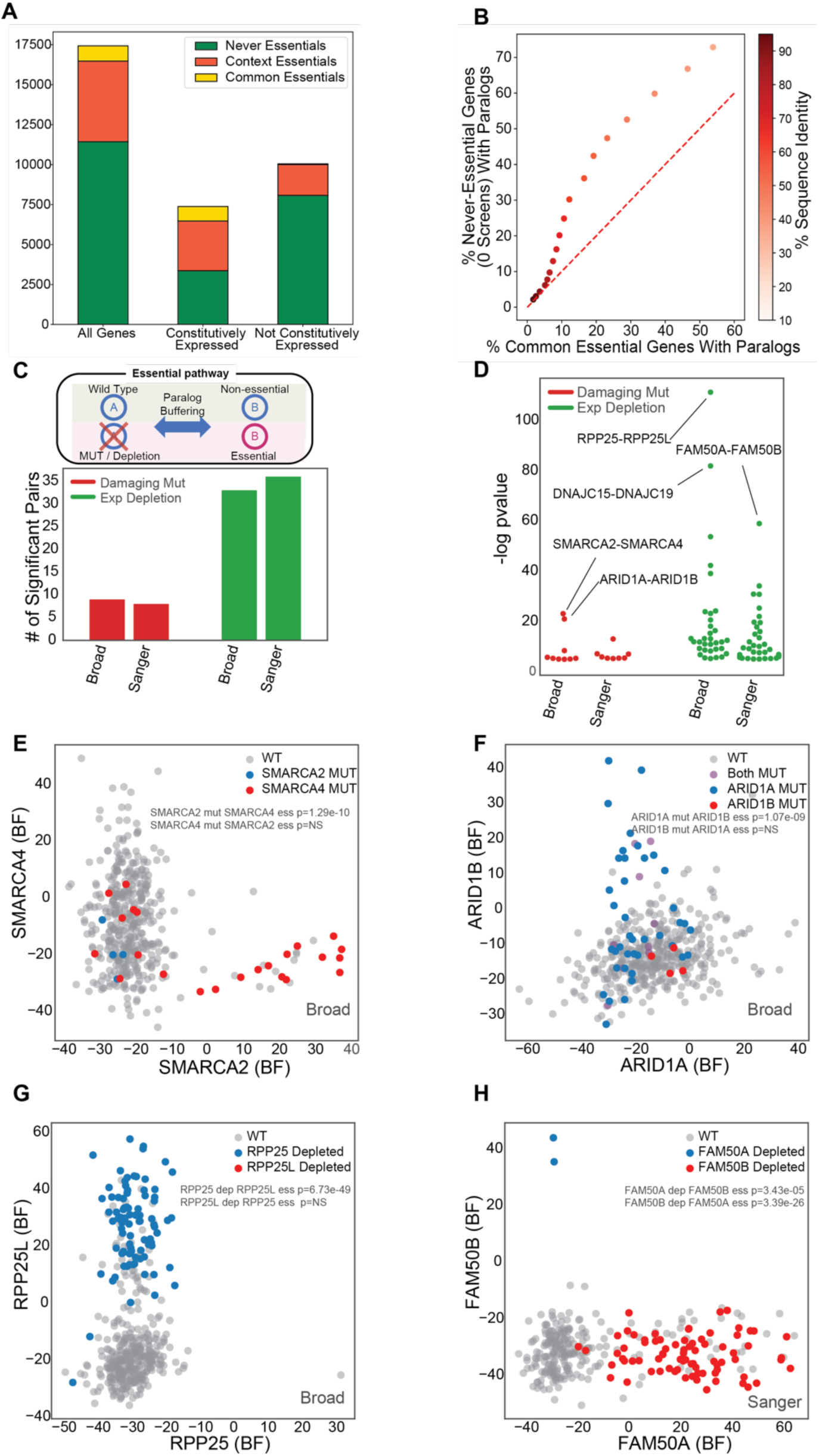
Paralogs are under-represented in CRISPR-Cas9 screens. **B)** Stacked bar graph showing the numbers of common, context and never essential genes among all genes, constitutively expressed and not constitutively expressed genes.Nearly half (46%) of all constitutively expressed genes are not observed as essential in any of the Avana screens. **B)** Scatter plot of the percentage of constitutively expressed never essentials with paralogs to that of constitutively expressed common essentials with paralogs based on shared percentage of sequence similarity thresholds. **C)** The number and **D)** significance of synthetic essentiality of paralog pairs identified in Broad and Sanger screens. Only pairs with p-value less than 0.05 were considered in these plots. **E-H)** Scatter plots of the synthetic lethality caused by mutational disruption of one paralog partner between **E)** SMARCA2-SMARCA4 **F)** ARID1A_ARID1B and by depletion of one paralog partner between **G)** RPP25-RPP25L and **H)** FAM50A-FAM50B.

These observations regarding the constitutively expressed genes raised the question about why we observe so few essential genes in these genetically heterogenous screens. Based on work in yeast (Hillenmeyer et al., 2008) and nematodes (Ramani et al., 2012), we naively assumed that all constitutively expressed genes should be essential in some context, and hypothesized that some combination of environmental or genetic buffering masks the fitness consequences of individual gene knockouts.

To examine the possible role of genetic buffering, we tested whether paralogs are overrepresented among the never-essentials. We obtained the list of the paralogs of human protein coding genes from Ensembl Biomart along with percent protein sequence similarity information (see Methods). After filtering for constitutively expressed genes, we observed paralogs show a wide range of amino acid sequence similarity, but that the majority of the paralog pairs as defined in Ensembl exhibit low similarity (Supplementary Figure 7B). To evaluate whether paralogs are enriched in never-essentials, we adopted a sliding scale of sequence identity and measured, at each threshold, the fraction of never-essentials and the fraction of common essentials captured. As shown in Figure 4B, as sequence similarity stringency is relaxed, never-essentials are much more likely to have a paralog than common essentials. At 45% or greater sequence similarity, nearly a third (30.2%) of constitutively expressed never-essentials have a paralog, compared with only 12.2% of common essentials, a ratio of ∼2.5:1. This is highly consistent with previous observations by De Kegel and Ryan (De Kegel and Ryan, 2019) and by Dandage and Landry (Dandage and Landry, 2019), who used similar approaches.

To further validate this observation, we explored the Avana data to find cases where loss of function of one member of a paralog pair resulted in increased dependency on the other. Unfortunately, the cell lines screened by CRISPR knockout libraries only contain LOF alleles of a fraction of the candidate paralogs, limiting this discovery avenue to a few dozen pairs (Figure 4C-D). Unlike previous approaches, limiting the search for functional redundancy to constitutively expressed genes excludes false positives arising from tissue-specific expression of paralog family members. Two well-described cases in the *BAF* (SWI/SNF) complex were immediately apparent: mutations in *SMARCA4* are strongly associated with dependency on paralog *SMARCA2* (P<10^-10^; Figure 4E), and mutations in *ARID1A* are associated with *ARID1B* dependency (P<10^-9^; Figure 4F). Expanding loss-of-function to include significantly depleted gene expression also reveals an emergent dependency on *RPP25L* when *RPP25* is depleted (P<10^-52^; Figure 4G). The two genes encode redundant subunits of RNAse P, a ribonuclease critical for maturation of tRNA, whose functional buffering was previously observed (Wang et al., 2015). A fourth example is *FAM50A/FAM50B* putative functional redundancy (Figure 4H). Interestingly, virtually nothing is known about the biological role of these genes.

## Discussion

CRISPR technology has revolutionized mammalian functional genomics and cancer targeting by leveraging endogenous DNA repair machinery to generate gene knockouts on a genomic scale. Extensive screening of cancer cell lines under the DepMap project—and, critically, the open availability of this data--affords an opportunity for re-evaluating the assumptions under which these assays have been carried out. Notably, assumptions about replication and library- and batch-specific effects have been addressed in some detail (Wang et al., 2019; Dempster et al., 2019; Rauscher et al., 2018), but questions about what might be systematically missing from these data have, to our knowledge, not been rigorously explored.

We developed a model that estimates the true positive rate, false positive rate, and false negative rate from a typical genome-scale CRISPR screen. The model which most closely matches data from a large panel of CRISPR screens suggests that a typical cell expresses 1,600-1,900 essential genes, but a single knockout screen only detects ∼80% of these, and multiple screens are required to saturate the essential genes of any tissue or tumor subtype. Hits among highly expressed genes are often replicated but false negatives are more prevalent among genes expressed at moderate levels. This carries severe implications for the identification of differentially essential genes, in particular using isogenic cell lines to identify synthetic lethals, as it suggests numerous replicates need to be screened in order to confidently discriminate cell-line-specific hits from false negatives/false positives.

A further implication of the false negative rate is that most cells/tissue types carry a larger number of overlapping essential genes that had been previously recognized. After allowing for false negatives, we identify nearly a thousand genes that are observed across all lineages that are deeply sampled in the Avana screens. In contrast, we find only 300-600 tissue-specific genes per lineage, with most showing overlap between related tissues. In fact, each tissue only carries at most a few dozen tissue-unique genes, and these are highly enriched for lineage-specific transcription factors.

Finally, we note that there are about 7,000 genes that are constitutively expressed in each cell, but only about half of these are ever detected as essential. Studies in model organisms suggest that virtually every gene shows a growth phenotype under some environmental condition (Hillenmeyer et al., 2008; Ramani et al., 2012). It is unknown whether this holds true for individual mammalian cells, though tumors are often modeled as though they are colonies of single-celled organisms. It is also the case that most genetic screens of tumor cells are carried out under permissive growth conditions, minimizing nutrient and oxidative stress to maximize growth rate and improve detection of dropouts. Thus, the degree of environmental buffering is largely unknown for these constitutively expressed never-essentials.

However, these never-essentials are highly enriched for paralogs. They are as much as 2.5 times more likely to have a paralog than always-essentials, suggesting that functional redundancy by related genes masks detection of a substantial population of genes in monogenic CRISPR knockout screens. This has profound implications for efforts to match targeted drugs with tumor genotypes, and to discover new candidate drug targets. Targeted small molecules often don’t discriminate, or discriminate poorly, between closely related paralogs, and it may be their promiscuity rather than their specificity that renders them effective. For example, MEK inhibitor trametinib effectively targets the protein products of both *MAP2K1* and *MAP2K2*, redundant kinases downstream of RAS/RAF oncogenes, but the functional redundancy of these genes renders them both invisible to monogenic CRISPR screens, even in RAS/RAF backgrounds (Kim et al., 2019). A systematic survey of paralogs families, in particular those with only two family members that are constitutively expressed, for synthetic lethality could unmask this genetic buffering and yield greater insight into the constellation of targetable genes in a tumor. All in all, systematic CRISPR screens represent an impressive first step toward characterizing tumor-specific genetic vulnerabilities, but genetic and environmental buffering may mask a substantial number of new targets.

## Methods

### Essentiality data generation

A raw read count file of CRISPR pooled library screens for 517 cell lines using Avana library (Broad DepMap project 18Q4) was downloaded from the data depository (https://figshare.com/articles/DepMap_Achilles_18Q4_public/7270880). Also, we downloaded Project Score (Sanger) screen (Behan et al., 2019) raw read counts for 323 cancer cells from the data depository (https://score.depmap.sanger.ac.uk/). We filtered the dataset to keep only the protein-coding genes for further analysis and updated their names using HGNC (Yates et al., 2017) and CCDS (Farrell et al., 2014) database. We discarded sgRNAs targeting multiple genes in Avana library to avoid genetic interaction effects. The raw read counts were processed with the CRISPRcleanR (Iorio et al., 2018) algorithm to correct for gene-independent fitness effects and calculate fold change. After that, the CRISPRcleanR processed fold changes of each cell line were analyzed through updated BAGEL v2 build 110 (https://github.com/hart-lab/bagel). In comparison with published BAGEL version v0.92 (Hart and Moffat, 2016), the updated version employed a linear regression model to interpolate outliers and 10-fold cross validation for data sampling. Essentiality of genes was measured as Bayes Factor (BF) based on gold standard reference sets of 681 core essential genes and 927 nonessential genes (Hart et al., 2014; Hart et al., 2017). Positive BF indicates essential genes and negative BF indicates non-essential genes. Lists of core essential genes and nonessential genes used in this study have been uploaded on the same repository with BAGEL v2 software. The screen quality was evaluated by using “precision-recall” function in BAGEL software, and F-measure (BF = 5), which is the harmonic mean of precision and recall at BF 5, was calculated for each screen. Finally, 446 cell lines for Broad screen and 320 cells for Sanger screen were selected for further study by F-measure threshold 0.8 to prevent noise from marginal quality of screens.

### Cumulative Essentials Analysis

A cumulative analysis approach was used to evaluate the cumulative distribution of essential genes and calculate the total number of true essentials (true positives), as well as the error rate (false discovery rate, FDR) in a screen. This is based on the principle described previously (Hart et al., 2014) that for screens with zero FDR, you would expect a cumulative essential gene observations plot that flattens out with a slope of zero at the total number of hits. For the actual screens, when the screens are repeated, the true hits are saturated but because false positives and cell-line specific essential genes might also be captured, you get a cumulative essentials curve with a positive slope as more false positives and context essentials are added. The BFs for all cell lines in the Avana18Q4 data were constructed in a matrix. We modeled the cumulative essentials plot in any sets of 20 screens from the Avana data. First, the initial set of 20 cell lines were sampled without replacement and the essential genes in the first screen out of 20 were identified with a BF of greater than or equal to 10. Then, the next screen was analyzed to obtain the essential genes in that screen and the newly discovered essential genes that were not present in the previous screen were added to the list of essential gene hits to obtain a cumulative essential gene list. The process was repeated for all subsequent cell lines to capture the cumulative essential genes in the 20 screens. The random sampling process was repeated for 100 times to sample different cell lines in different orders and prevent bias in the analysis. The resulting mean cumulative essentials curves were plotted with standard deviations of cumulative essential gene observations observed in each of 100 iterations. For each iteration, the newly discovered essential genes in each set of 20 screens were also identified and these new hits were also plotted on the same plot.

### Synthetic Genome Model-Estimating the number of essential genes per screen and screen average FDR

To estimate the number of essential genes per screen and the average error rate, we conducted in silico simulations of synthetic screens. A synthetic genome was constructed with the number of genes assayed in the Avana library (N=17427). For a given genome with N number of genes in it, there is a number of true essential genes; represented by n, and number of non-essential genes (N-n). Then, the precision (1 - False Discovery Rate) of the assay is represented by the ratio of true positives to that of the total number of hits. We defined a range of thresholds for false discovery rate to test in our model from 1-15%. We then randomly sampled this synthetic genome with a screen with randomly drawn 1267 hits (the mean number of essential gene hits at BF>10 across all Avana screens) from the essential (n) and non-essential (N-n) populations based on the defined FDR in the simulation (e.g. for 10% FDR, 127 nonessentials and 1,140 essentials were randomly selected). We observed the cumulative essential genes across 8 iterations since we estimated that sampling 8 screens was a good estimate of observing the trend in screen saturation in the Avana data. At the same time, we constructed a mean cumulative essentials curve determined from the Avana data for any 8 screens using bootstrapping for 100 iterations. After running the simulations for a range of different number of essential genes (n) and FDR values, cumulative observation curves were plotted for each simulation. The root mean squared deviation for every synthetic screen was calculated by evaluating the difference between the observed values and the cumulative essentials curve determined from the Avana data using the sklearn.metrics module in SciKits package for Python version 3.6.

Since the Avana data is composed of multiple different tissue types, it is possible that some of the tissue specific essential genes would be wrongly included in the false positives. To minimize this effect, we repeated the synthetic genome model in individual tissue types in the Avana data that were represented by at least 16 high-quality cell lines using the same parameters described above.

### Saturation Modeling-Modeling the number of high confidence essential genes and false positives

While the in-silico simulations enabled an estimation of the average number of essential genes per screen, they didn’t give information about which genes were truly essential. To distinguish essential genes from false positives, we identified the tissue types in the Avana data that were well represented (n>=16 screens). We concluded that 15 tissue types fit our criteria and for these tissues, we evaluated the frequency of essential gene observations in any 8 screens. For each tissue type, we randomly selected a set of 8 initial screens and identified the cumulative essential genes in this set and also plotted the frequency of essentiality observations. For our model, we assumed that if these first 8 screens have reached saturation (to approximate for our model), then the subsequent hits in the next set of screens would give us false positives. Therefore, we randomly selected a subsequent set of 8 screens from the same tissue type without replacement and looked at the newly discovered cumulative essential genes that were not present in the initial screens to model the frequency distribution of the false positives. This process was repeated 100 times for each tissue type and the resulting distribution of essential gene counts in these screens were plotted, which were used to calculate the bin-wise FDR. For the 100 iterations that were performed per tissue type, genes observed as essential in at least 3 screens on an average out of 8 were considered as high confidence essential in that tissue. Finally, we assessed how many genes were captured as essential in common in all tissues to find the set of “common” essential genes (n=954) and context essentials in each tissue type. We used the UpsetR package in R to visualize the set of intersections of essential genes in 15 tissue types.

### Expression data

We utilized the log2 transformed RNA-seq TPM expression data from Depmap Data Portal (https://depmap.org/portal/download/CCLE_depMap_18Q4_TPM_v2.csv). We filtered the TMP expression data for the tissues and cell lines used in our analysis. 393 out of 396 cell lines used in downstream analysis had corresponding expression data.

### Expression analysis

Mean TPM expression was calculated for all the genes in each of the bins from the histogram of the frequency of essentiality observations of the initial set of 8 screens representing the true positives. The expression values were plotted on a secondary y-axis with the error bars representing the standard deviation of TPM expression of the genes in each bin.

### Essentiality score distribution of genes in non-essential screens

To evaluate the mean essentiality score of genes representing true positives in the screens they were not observed as essential in, for each iteration, we determined the screens that the genes in each bin had a BF <10 and calculated a mean essentiality score for each gene in those screens.

We visualized the distribution of mean essentiality score observations of genes in each bin in the non-essential screens as a violin plot.

### Process enrichment analysis

From our modeling of the number of essential genes and false positives per tissue type, we had evaluated the frequency of essential gene observations in the 100 iterations we performed. We constructed a table of mean number of screen essentiality observations out of 8 screens, in 100 iterations for every gene in each tissue type. To investigate the trends of enrichment of essential pathways, we measured the functional enrichment of gene annotations depending on the thresholds for the mean number of screens that a gene was observed as essential in in 0.5 increments. For each threshold, we used the python gseapy package version 0.9.13 for python 3.6 to perform process enrichment in several databses including KEGG, CORUM and GO Biological Process using all 17,427 genes assayed in the Avana library as the set of background genes. Going in 0.5 increments in the frequency of screen essentiality observations out of 8 screens, we plotted the P-value of annotation enrichment for several terms to show the different trends in the data. For every gene in the processes that were enriched, we calculated their mean TPM expression in the corresponding cell lines that the process enrichment analysis was conducted. Finally, we evaluated the essentiality scores of the genes in the enriched pathways in the corresponding cell lines where the mean number of screen essentiality observation was greater than 0 out of 8 screens. We used the seaborn.clustermap function of the seaborn package to plot the hierarchically-clustered heatmap of the essentialty scores of the genes in each pathway in the corresponding tissue type using the average linkage method and the Euclidean distance metric.

### Defining constitutively expressed genes

The standard deviation of expression versus the mean expression values for all genes assayed in the Avana library (N=17,427) across all 393 cell lines, for which the expression data was available, were plotted. Constitutively expressed genes were defined as genes with mean expression > 2 log2(TPM) and standard deviation of expression expression <1 log2(TPM). With these thresholds we identified 7,378 constitutively expressed genes in the Avana dataset. Among these, 910 were commonly essential across all cell lines, 3107 context essentials showed essentiality profile in some of the Avana screens, whereas 3361 genes were not observed as essential in any screen.

### Paralogs

The human paralogous gene pairs for the protein coding genes were utilized from Ensemble Release 95 Biomart with GRCh38.p12 genome assembly (Zerbino et al., 2018). This release of Ensemble estimates paralogues from gene trees that are constructed with HMM as described in more detail at http://www.ensembl.org/info/genome/compara/homology_method.html. Other information such as chromosome location, paralogue percent sequence identity to human target gene and percent sequence identity of target gene to the paralogous gene were also downloaded. After removing duplicate gene pairs and filtering for constitutively expressed genes, the percent sequence identities of all human target genes to their paralogs were plotted against the percent sequence identities of the paralogs to the target human gene to reveal that the majority of the human paralogous gene pairs had low percentage sequence similarity. Next, the paralog pairs were binned according to different thresholds for percent sequence identity from a range of 10-95% and for each bin the percentage of constitutively expressed never essentials with paralogs and the percentage of common essentials with paralogs were calculated and their distributions were plotted.

### Investigation of evidences of functional redundancy between paralog genes in CRISPR screens

To investigate the functional redundancy between paralog genes in Broad and Sanger screens, we conducted statistical test of synthetic essentiality which is defined that a gene is essential when the other paralog partner is disrupted by damaging mutations (frameshift or nonsense) and depletion of expression (mean log TPM < 1.0). Statistical significance was calculated for all one-one paralog pairs with at least 30% sequence similarity by Fisher’s exact test with a contingency table of presence of disruption in gene A and binary essentiality of gene B (BF threshold = 10). We found total 9 and 33 synthetic essentiality paralog pairs from Broad screens by mutation evidences and expression depletion evidences, respectively. Also, we found total 8 and 36 synthetic essentiality paralog pairs from Sanger screens by mutation evidences and expression depletion evidences, respectively.

### Essential gene comparisons in different datasets

Previously defined core essentials from (Hart et al., 2017) and (Behan et al., 2019) were downloaded from their corresponding supplementary data sections. Raw read count data from Sanger Institution’s Project Score (Behan et al., 2019) was downloaded and processed with the same pipeline using CrisprCleanR and BagelV2 algorithms to generate Bayes factors for each gene in every screen. Genes located on the X-chromosome were removed from the core essential gene set V2 (CoreV2, from (Hart et al., 2017)) and Sanger data (Behan et al., 2019) prior to comparisons. The common cell lines between Avana and Sanger screens that have F-measures of greater than or equal to 0.8 were identified (N=117 cell lines). After filtering for genes assayed in both libraries, common essential genes and unique hits specific to each dataset were investigated. For Sanger specific core essential genes, their Bayes Factors in the common cell lines between Sanger data and Avana data were compared and gseapy module of Python3.6 was used to perform biological process enrichment using ‘GO_Biological_Process_2018’ gene set.

## Supplementary Figures

**Supplemental Figure 1.**
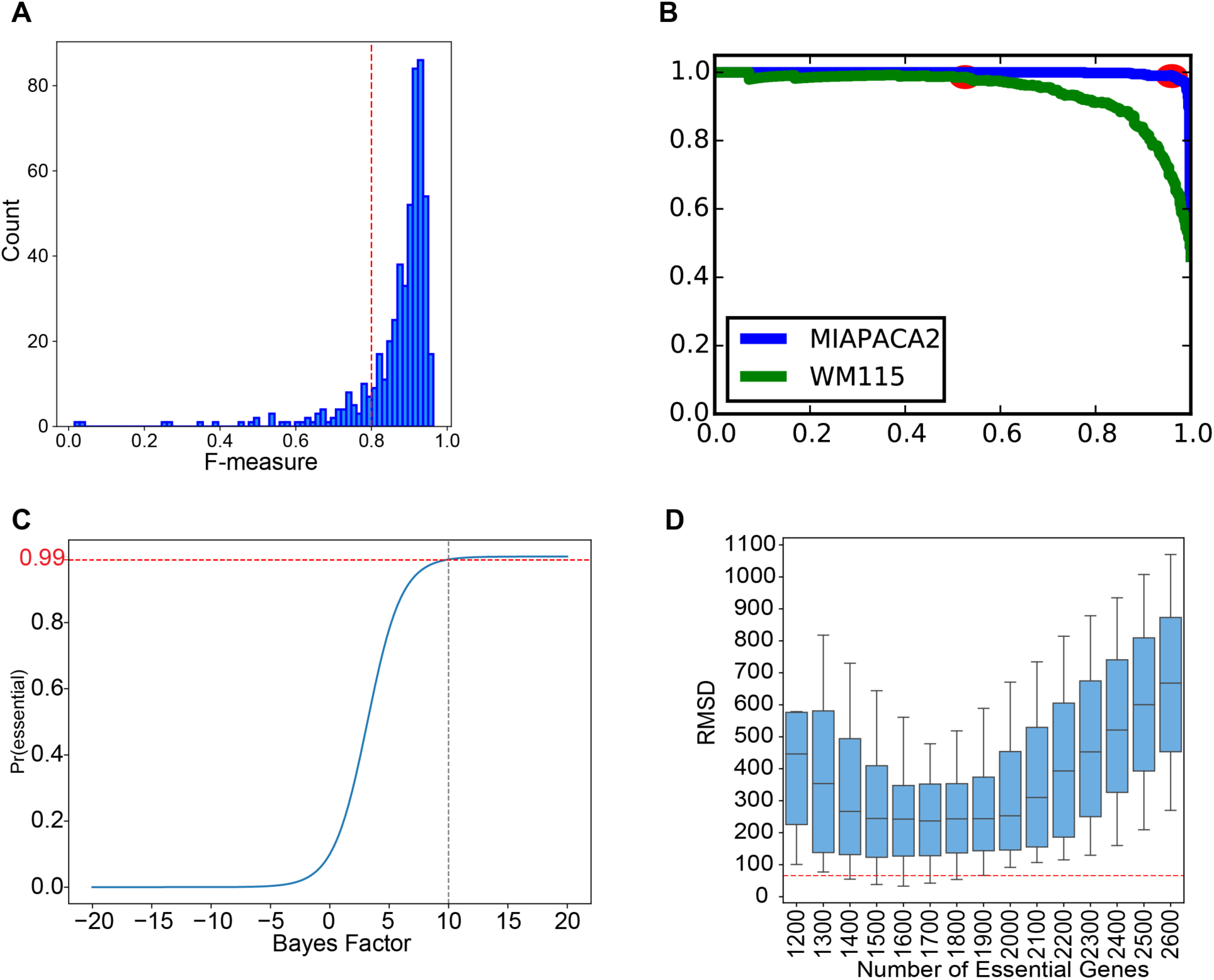
**A)** the distribution of the F-measures of 517 Avana Screens. The red dashed line indicates F-measure of 0.8 and screens with F-measure >0.8 were considered as high-performing and were retained for downstream analyses. **B)** Precision-recall curve specifying an example of a good performing screen (MIAPACA2, in blue) and a bad performing screen (WM115, in green). For all Avana screens, precision-recall curves were calculated using the reference gold standard sets of essential and non-essential genes and the point on the precision-recall curve for each screen where the BF crossed 5 (red points) were identified and the F-measure of each screen was calculated at that point. **C)** Bayes factor of 10 represents a strict threshold corresponding to a posterior probability of gene essentiality of ∼99% **D)** The distribution of the root mean squared deviation (RMSD) values for each simulation reveals a range of models with RMSD < 2xRMSD_min_ indicated by the red dashed line.

**Supplemental Figure 2.**
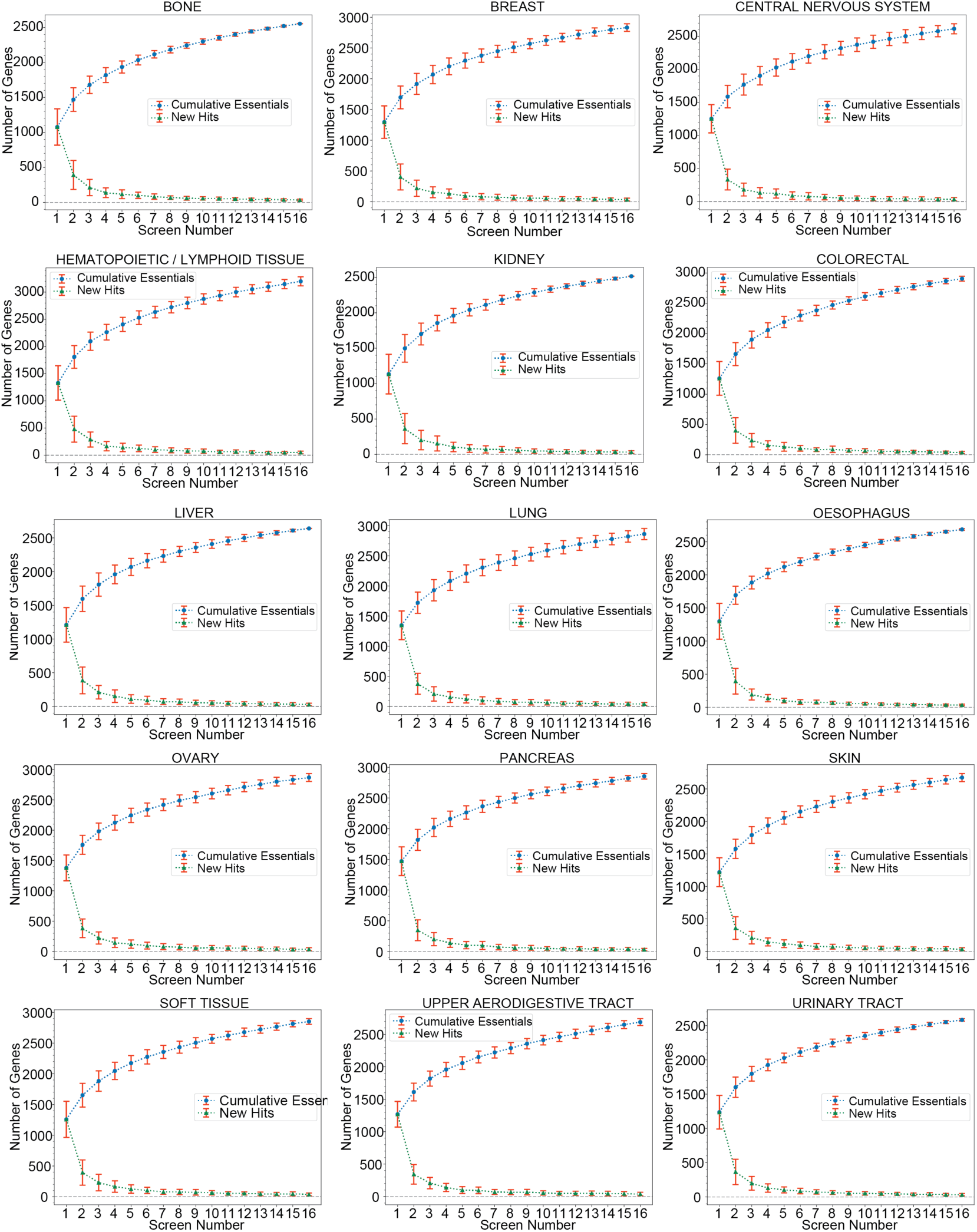
The cumulative essentials curves for lineages represented by more than or equal to 16 high-quality screens. Sets of 16 cell lines were randomly selected without replacement from all screens and the number of cumulative essential genes with BF>=10 in each consecutive screen were plotted in blue with the error bars indicating the standard deviation of cumulative essential gene observations across 100 iterations. The number of newly discovered essential genes in each consecutive screen was also plotted in green.

**Supplemental Figure 3.**
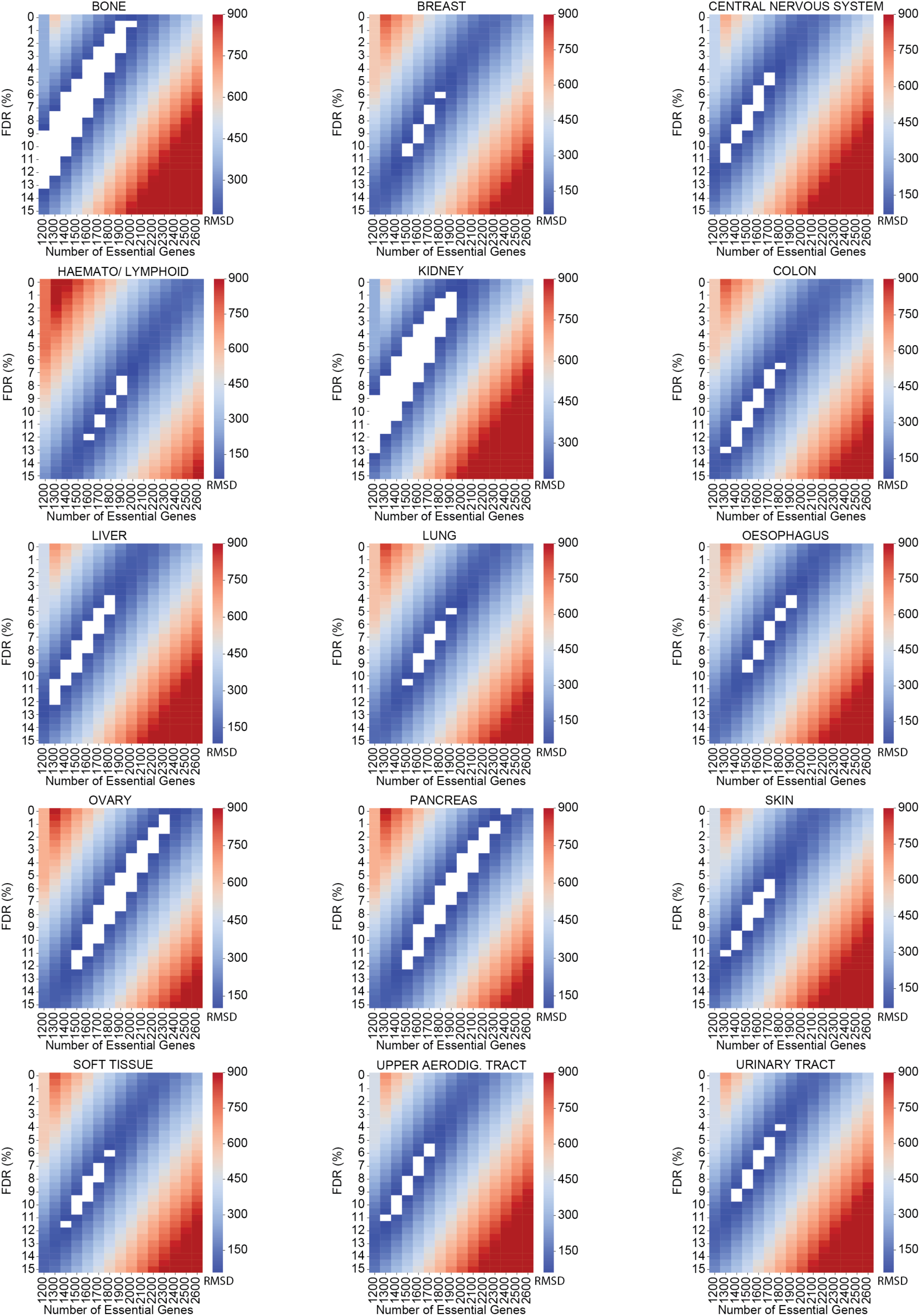
Synthetic genome modeling, applied to each tissue type, estimates the number of essential genes and false discovery rate (FDR) per tissue. Heatmaps showing the root mean squared deviation (RMSD) for the models versus the FDR and the number of essential genes in each simulation. The white boxes indicate models with RMSD <2 x RMSD_min_.

**Supplemental Figure 4.**
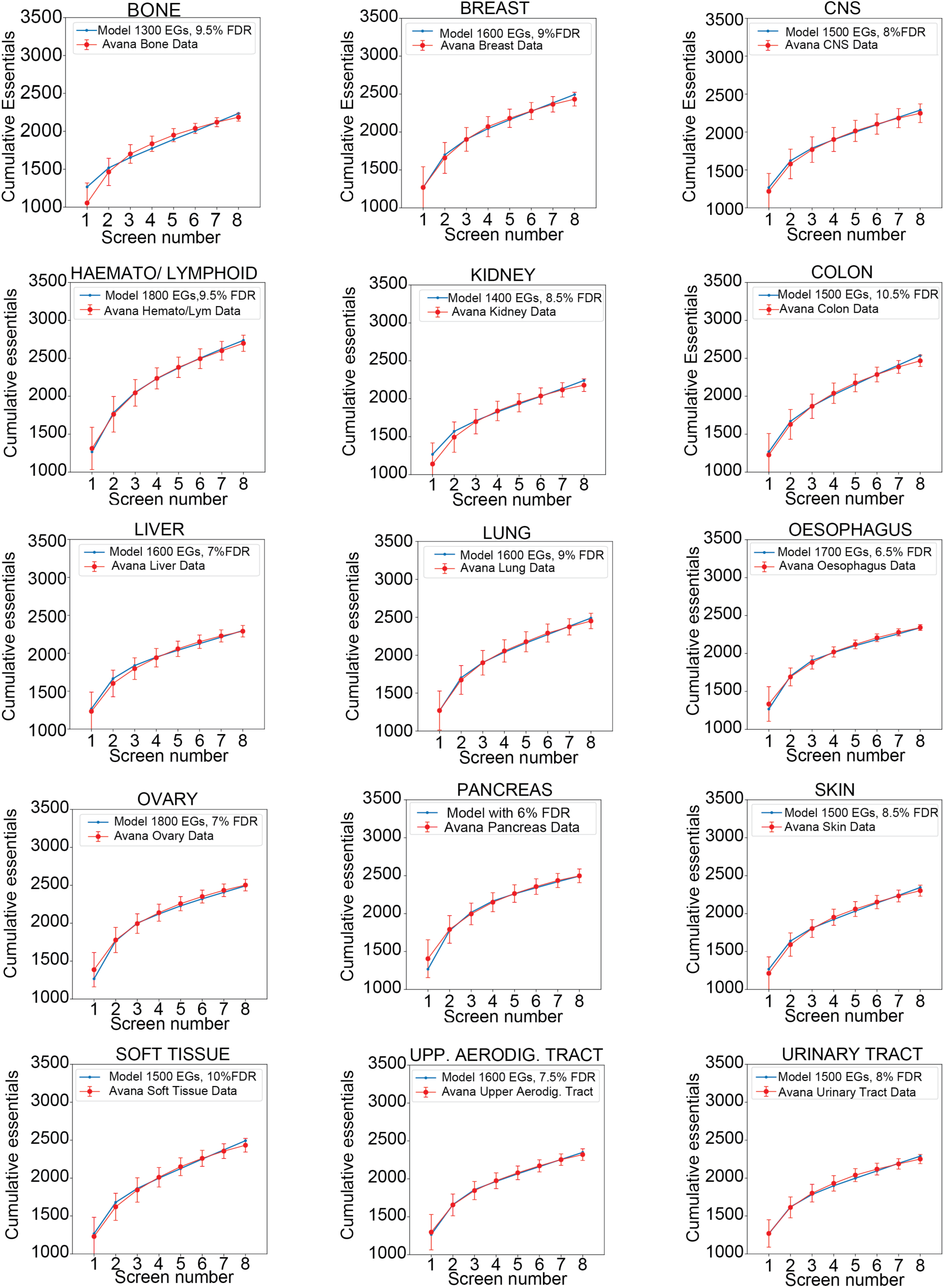
The best fitting models from the synthetic genome modeling approach for individual tissue types. The cumulative essentials curves were plotted for the best fitting model indicated by the blue lines and their fit to the Avana data in their corresponding tissue types (cumulative essential genes across sets of 8 call lines randomly selected without replacement from all available screens in that tissue type for 100 iterations) is shown in red.

**Supplemental Figure 5.**
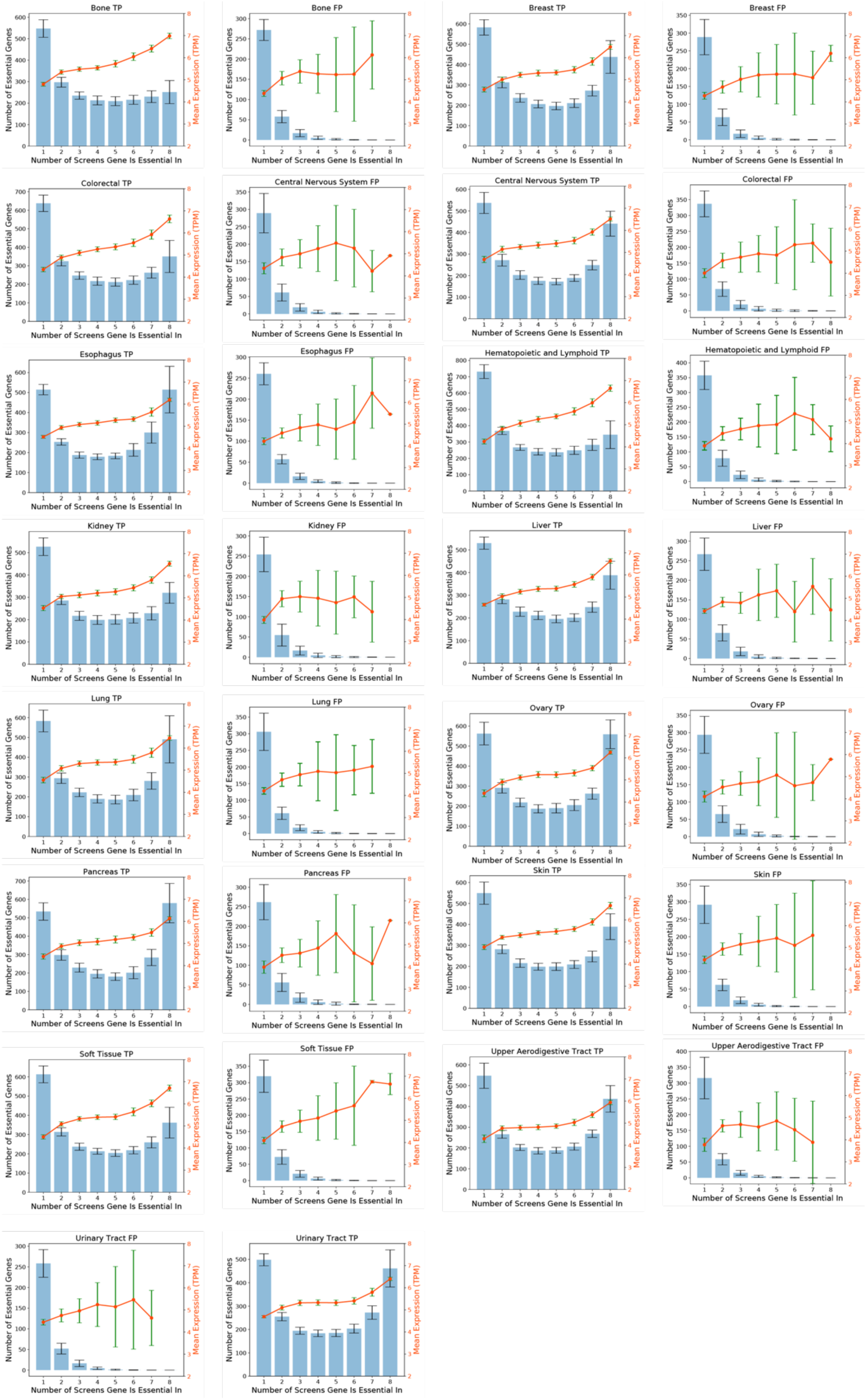
The number of genes in each bin and the mean mRNA expression (TPM) of the genes (indicated by the secondary Y-axis in orange) in corresponding bins for each tissue type for putative true positives (TPs) and false positives (FPs). Error bars indicate the standard deviation of expression of genes in each bin.

**Supplemental Figure 6.**
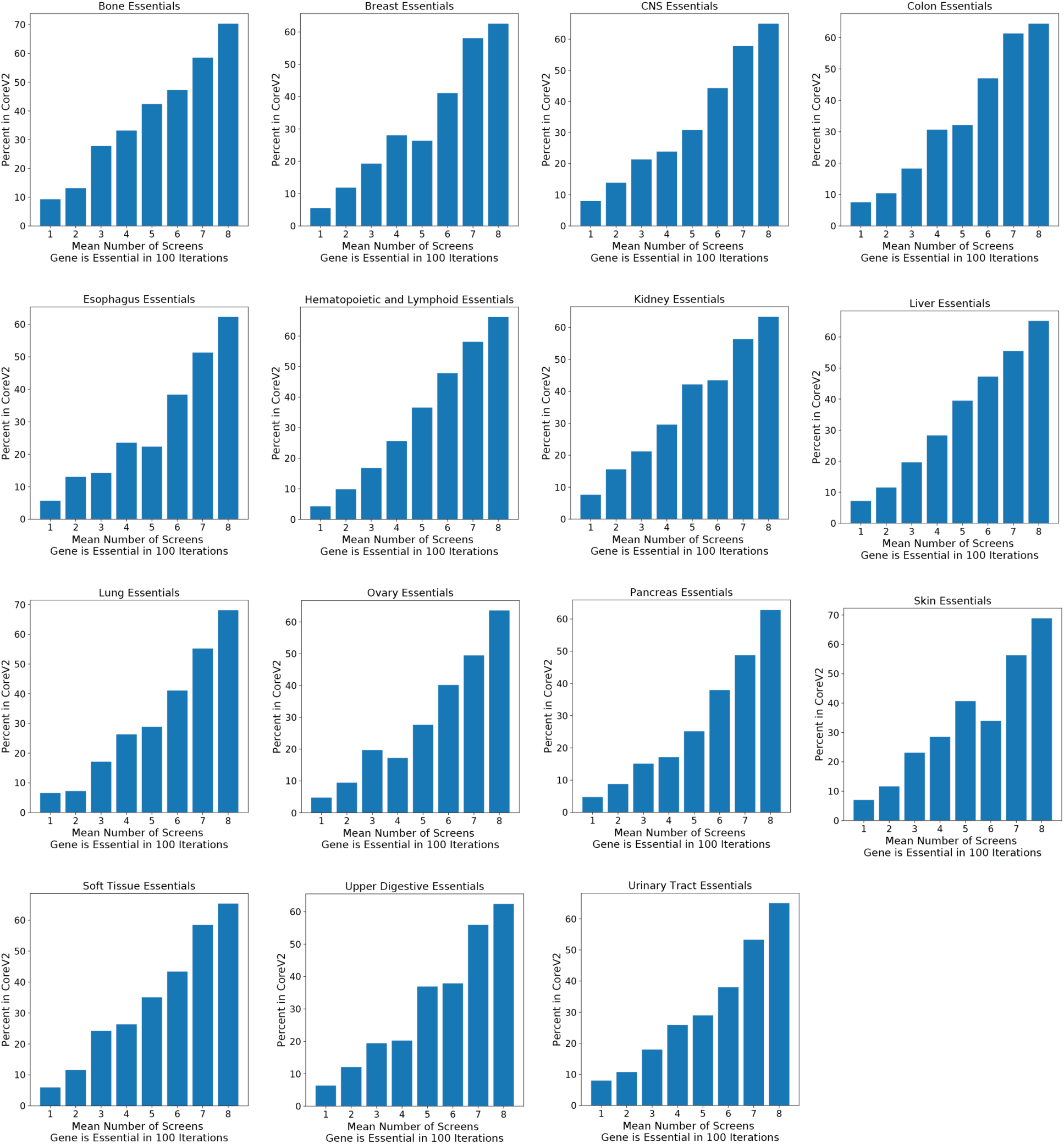
Bar plots in each tissue type showing, for genes binned according to the mean number of screens they are observed as essential in 100 iterations, the distribution of their percentage in the previously defined gold standard set of core essential genes (Core V2).

**Supplemental Figure 7.**
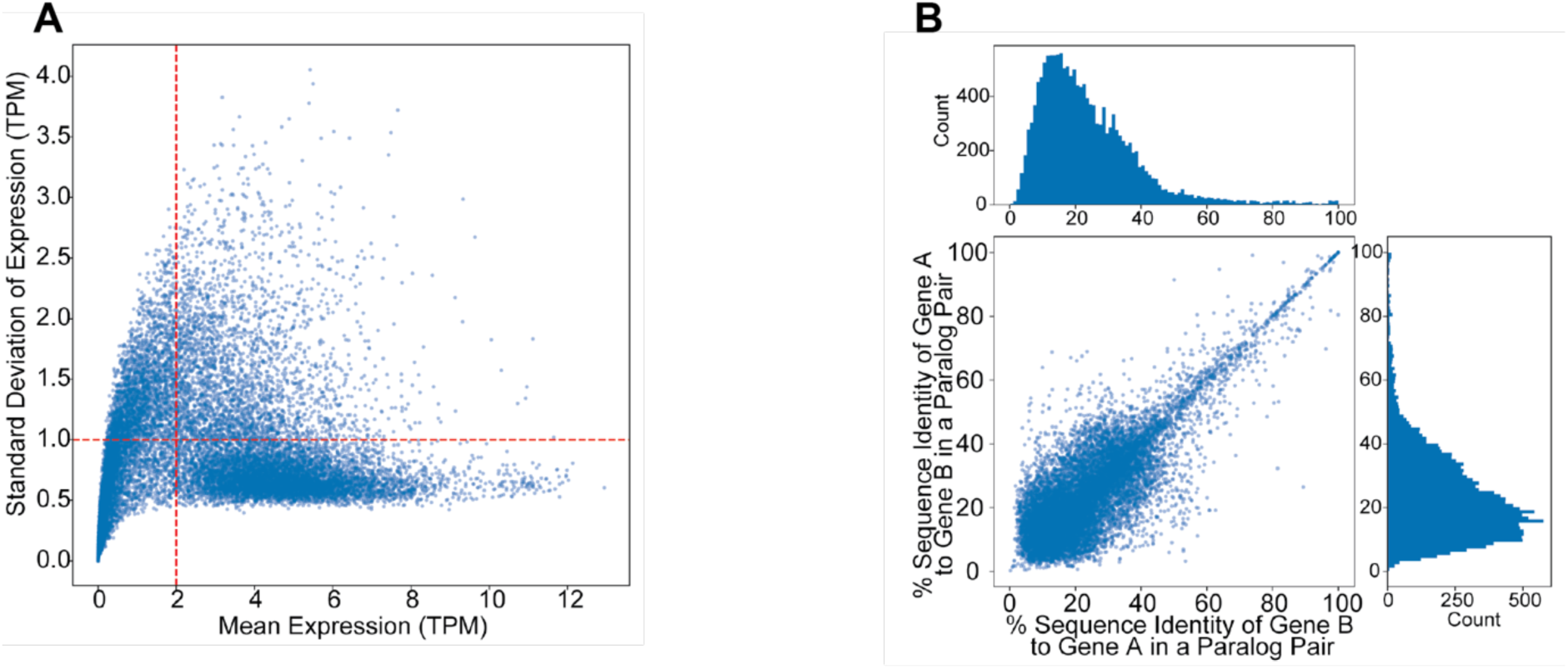
**A)** Scatter plot of the standard deviation versus mean mRNA expression (TPM) of all genes assayed in the Avana data across all cell lines for which the expression data was available. The red dashed lines indicate the thresholds to define constitutitvely expressed genes with mean logTPM>2 and stdev <1 (n=7,378 genes). **B)** Scatter plot and marginal histograms of the constitutively expressed paralogs in Ensemble have low amino-acid sequence similarity.

## Supplementary Table Legends

**Supplementary Table 1.** Bayes Factors for the 446 cell lines used in this study with F-measure above 0.80 post CrisprCleanR processing.

**Supplementary Table 2.** Table of binwise false discovery rates across 100 iterations for the tissue types investigated in this study.

**Supplementary Table 3.** Gene frequency observations out of 8 screens across 100 iterations by tissue type.

**Supplementary Table 4.** Table of 954 common essential genes and high confidence context essential genes in each tissue type.

**Supplementary Table 5.** Common essential and core essential genes unique to each approach among previously defined core essential genes.

## Acknowledgments

MD and EK were supported by the Cancer Prevention Research Institute of Texas (CPRIT) grant RR160032. TH is a CPRIT Scholar in Cancer Research, and is supported by NIGMS grant R35GM130119 and MD Anderson Cancer Center Support Grant P30 CA016672.

